# Inclusive fitness forces of selection in an age-structured population

**DOI:** 10.1101/2022.06.02.494506

**Authors:** Mark Roper, Jonathan P. Green, Roberto Salguero-Gómez, Michael B. Bonsall

**Author notes:** Corresponding author; Mark Roper **Email:**.

## Abstract

Current evolutionary theories of senescence predict that the force of selection on survival will decline from maturity to zero at the age of last reproduction, and the force of selection on reproduction will decline monotonically from birth. These predictions rest upon the assumption that individuals within a population do not interact with one another. This assumption, however, is violated in social species, where an individual’s survival and/or reproduction may shape the fitness of other group members. In such species, it is inclusive fitness that natural selection optimises. Yet, it remains unclear how the forces of selection on survival and reproduction might be modified when inclusive fitness, rather than population growth rate, is considered the appropriate metric for fitness. Here, we derive inclusive fitness forces of selection for hypothetical populations of social species. We show that selection on survival is not always constant before maturity, and can remain above zero in post-reproductive age classes, contrary to conventional models of senescence. We also show how the trajectory of the force of selection on reproduction does not always decline monotonically from birth, as predicted by classical theory, but instead depends on the balance of benefits to direct fitness and costs to indirect fitness. Our theoretical framework provides the unique opportunity to expand our understanding of senescence across social species, with important implications to species with variable life histories.

## Main Text

To date, there are no general theories for how senescence might evolve differently in groups of social species. At the demographic level, senescence is defined as the decline in organismal fitness with increasing age (1). Hamilton (2) provided a mathematical explanation for the seemingly counter-intuitive evolution of senescence: the force of natural selection weakens with age, and so detrimental alleles acting late in life can persist despite their negative effects on fitness (3 – 6). Two years prior, Hamilton (7 – 8) also introduced the concept of inclusive fitness, which has had a profound impact on our understanding of the evolution of social life histories (9 – 11). Inclusive fitness quantifies *(i)* an individual’s number of offspring in the absence of social effects and *(ii)* the effects an individual has on the number of offspring produced by other individuals, weighted by relatedness (7 – 8). It has not yet been fully considered from a theoretical standpoint, however, how these effects an individual has on the fitness of others may alter the evolution of senescence.

An age-specific force of selection describes the relative effect on fitness at different age classes of a mutant allele that impacts survival or reproduction. How might the components that contribute to such age-specific forces of selection differ between a solitary and a social species? First, consider an individual of a solitary species. When this individual dies, it loses access to any future reproduction it might have achieved. If a mutant allele arises in this population that increases the risk of dying at a certain age, say *x*, then the force of selection that acts against the allele is proportional to the expectation of residual reproduction that the individual may have realised (2). Now, imagine instead a social species in which individuals within a group influence one another’s survival and reproduction, for example, through the provision of alloparental care or through competition for limiting resources. For an individual, death means the loss of any future reproduction, just as in the solitary case. However, in social species, an individual’s death may also alter the survival and reproduction of other individuals (12 – 13). For instance, the death of an individual providing alloparental care may lead to a reduction in breeder productivity. Alternatively, where there is competition within groups for resources, the death of an individual may release resources that other group members may use for survival and reproduction. If individuals within a group are related, then these effects will be under kin selection. For example, an increase in mortality late in life can be adaptive if relatives stand to benefit from the death of a focal individual (14 – 20). On the other hand, mortality may be more strongly selected against if individuals can transfer beneficial resources to others (21 – 23). When the death and reproduction of a focal individual not only impacts its own fitness, but also the fitness of relatives, the force of selection acting on a mutant allele at age *x* must also consider these complex social effects.

To incorporate social interactions into the evolutionary theory of senescence, we develop a general model for quantifying age-specific inclusive fitness forces of selection in social species. Here, we focus on the effects of cooperative interactions between individuals and the corresponding forces of selection, but note that our model also has scope to consider other scenarios, such as cases of harm (see **Discussion**). Using an infinite island framework to describe a resident social population (16, 20, 24 – 34), we explore the fate of a mutant allele that alters *(i)* survival rate from age *x* to age *x* + 1 and *(ii)* reproduction at age *x*. We derive inclusive fitness forces of selection acting on these mutant alleles, which indicate how the efficacy of natural selection changes with age with respect to socio-demographic parameters. After deriving general analytical results, we explore the applicability of our framework to different social settings by providing numerical solutions for two examples of social structures: *(i)* the grandmother hypothesis: post-reproductive individuals aiding juvenile survival and *(ii)* cooperative breeding: juveniles aiding reproduction by adults. We conclude by discussing the implications and possible extensions for our model.

## Model

We consider a population divided into an infinite number of patches, and model the population dynamics of a focal patch. This infinite island approach (16, 20, 24 – 34) allows kin selection to be modelled while also considering the effects of demography, which is appropriate for considering an age-structured population in which individuals have effects on one another’s fitness. Each patch, which could also be conceptualised as a territory, contains discrete groups of exactly *N* individuals that are, for simplicity, haploid and asexual. We also assume that patches produce a large number of offspring in each breeding season so that no position on any patch is vacant at the start of each breeding season (*i*.*e*. a density-dependent stationary population). Offspring that establish on to a patch are designated age 1 and can survive until some maximum age, *ω*, at which point they die. Time proceeds in a series of discrete breeding seasons, during which individuals have a probability of surviving to the next breeding season, *p*(*x*), and a rate of reproduction, *b*(*x*), that may vary with age, and can be described by a population matrix model (***A***). Individuals may receive contributions to their survival and reproduction from the other *N* − 1 individuals on their patch, and may themselves contribute to the survival and reproduction of the *N* − 1 conspecifics on the patch.

Fundamental to this model is the concept of ‘transfers’. Biologically, transfers represent the help or harm to other individual’s fitness components: survival and reproduction. Transfers occur in the currency of genetic offspring equivalents, the same currency as survival and reproduction. Here, we assume that the transfers an individual makes to others is a function of the ages of both the actor and recipient (Fig. 1). We display transfers between individuals as 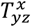: if *y* = 1, this represents an individual in age class *x*’s social effect on the reproduction of age class *z*, while *y* = *z* + 1 would represent an individual in age class *x*’s social effect on the survival of age class *z*.

**Figure 1.**
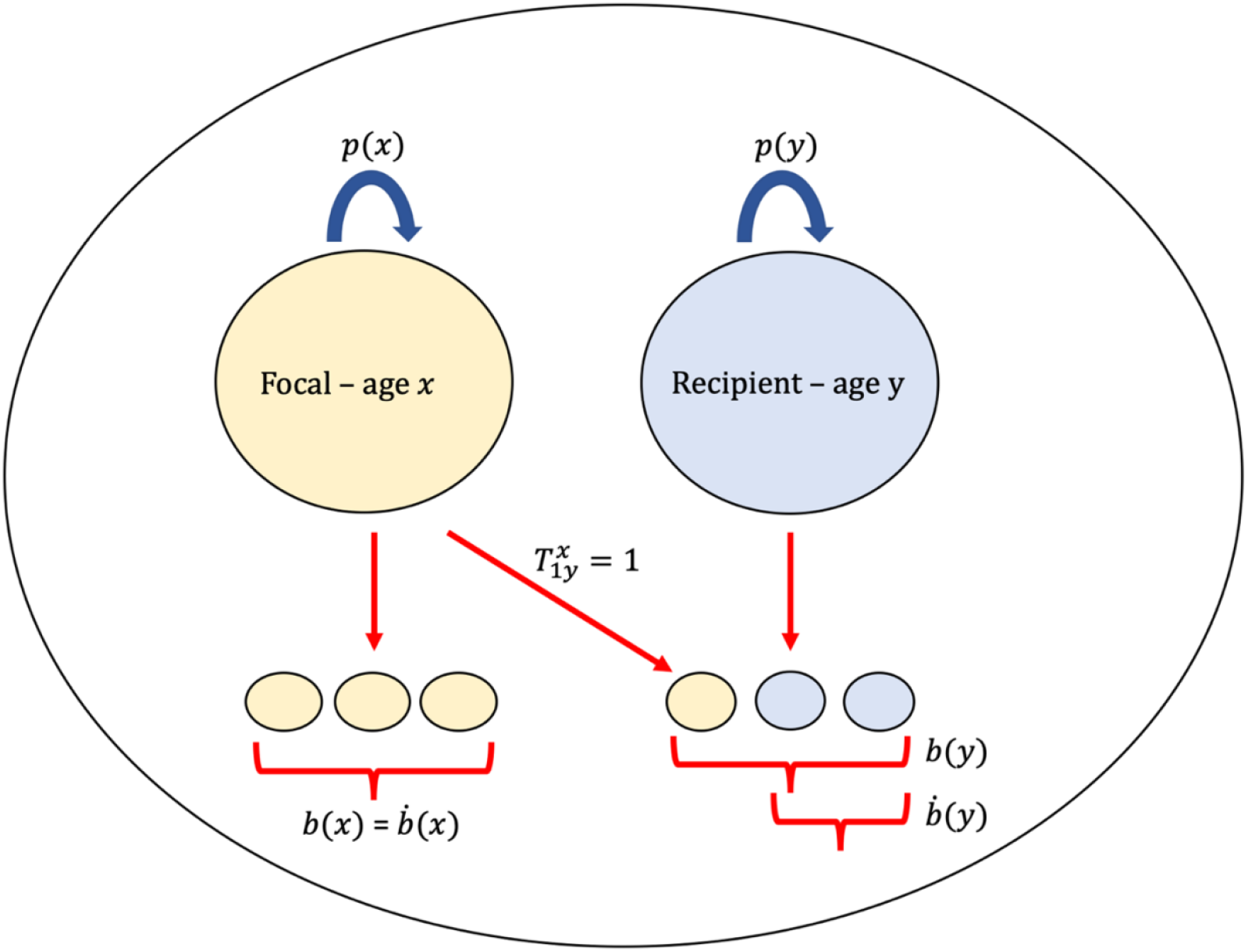
An example of a genetic offspring transfer between two individuals using inclusive fitness. To illustrate transfers, we consider a patch with two individuals, one of age *x* and the other of age *y*. The individual aged *x* has *b*(*x*) offspring, survives with probability *p*(*x*), and receives no social transfers from other individuals in the population when aged *x*. We imagine a social behaviour exists whereby the individual aged *x* contributes to the reproduction of individuals aged *y*. In this scenario, the individual aged *y* has *b*(*y*) offspring in the current breeding season, but one of these offspring is due to the transfer from the focal individual aged *x*. Following inclusive fitness logic, the offspring produced due to the social behaviour of the individual aged *x* is stripped from the inclusive fitness of the individual aged *y*, leaving 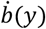 as their inclusive fitness contribution to age class 1. The inclusive fitness contribution of the focal individual aged *x* to age class 1 is 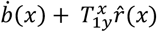, where 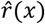 represents the relatedness of an individual aged *x* to the offspring it helped to produce.

To quantify the inclusive fitness contributions of a focal individual of age *x*, a series of key considerations must be made. Specifically, we must *(i)* exclude the fraction of the class-*y* offspring of a focal class-*x* individual that are born or survive as a consequence of the social environment (the help or harm of other individuals), and *(ii)* augment the total production of class-*y* offspring from all other age classes, including other individuals in age class *x*, that are born or survive due to the social contributions of a focal class-*x* individual. These latter offspring contributions are weighted by the coefficient of relatedness between an individual of age class *x* and the class-*y* offspring of the recipient class (7 – 8). For example, a focal individual aged *x* survives with probability *p*(*x*) and has a rate of reproduction *b*(*x*). A fraction of these rates of survival and reproduction may be due to social interactions. These fractions are excluded from the inclusive fitness of the focal individual, leaving 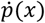 and 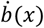, with dot notation representing the effect of a focal individual’s own genotype on its own survival or rate of reproduction, *i*.*e*. direct fitness. Of the 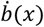 offspring produced due to the genotype of an individual aged *x*, a proportion *d* disperse, and a proportion 1 − *d* remain at their natal patch. A fraction *c* of the dispersing offspring die, representing a cost of dispersal. Surviving, dispersed offspring are evenly distributed among all sites, regardless of distance, and compete (fair lottery) for sites freed by adults that die in the current breeding season. Asymmetric competition is assumed so that juveniles do not displace resident adults, and die if they do not gain a breeding position on a patch. Offspring of a focal individual aged *x* face a probability of establishment *g*(*x*) onto their natal patch, and 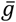 on a different, random patch in the population.

In a population with social interactions between patch members, we can populate a matrix (***W***) with the inclusive fitness (genetic offspring) contributions of individuals in age class *x* to individuals in age class *y* (*w*_*yx*_):

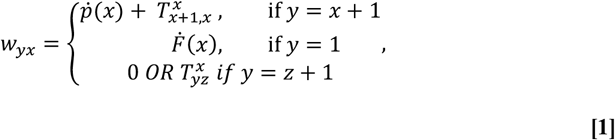

where

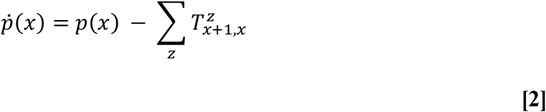

and

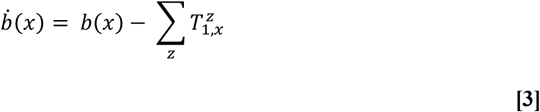

and

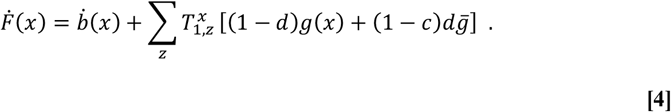

The proportions of the survival and reproduction of a focal individual aged *x* that are due to the genotypes of other individuals are represented in the summation terms on the right-hand side of **[2]** (survival) and **[3]** (reproduction) (where 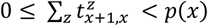, and 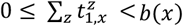. Importantly, these proportions are distributed to other age classes, thus ensuring that no offspring is ‘double counted’ (35 – 36). A focal individual of age *x* may also contribute to the survival and reproduction of others, accumulating indirect fitness through the transfer of genetic offspring. Contributions to survival are captured as 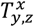 (where *y* = *z* + 1, and *y* ≠ 1), and reproduction as 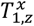(summed across age classes to equal 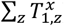). The magnitude of these contributions will depend on i) the expected number of individuals in the recipient age class, ii) the fraction of the total contribution of all age classes combined to the survival or reproduction of the recipient age class individuals that is due to a focal individual aged *x*, and iii) the relatedness between a focal individual aged *x* and an individual in the recipient age class (see **Supplementary Information Appendix C**). This approach to modelling social interactions assumes that there are fractions of survival and fecundity of each age class that are due to the social environment (which could equal zero), and that these fractions are distributed to other individuals across age classes. If there are no explicit social interactions between multiple individuals on a patch, equation **[2]** simplifies to a population with limited dispersal and Ronce & Promislow’s (20) kin competition selection gradients can be computed. With full dispersal (no offspring stay at the patch in which they’re born) and no social interactions, equation **[1]** simplifies to Hamilton’s panmictic population, and his forces of selection can be computed (2).

### An inclusive fitness force of selection

To compute forces of selection, we are ultimately concerned with a hypothetical mutation that alters survival rate or rate of reproduction at age *x*. The derivative of the growth rate of the mutant population, λ, with respect to the phenotypic effect of the mutation, δ, gives an indicator of the force of selection acting on the mutant allele (2, 20, 37 – 38). We consider mutations of weak effects (small δ) and first-order effects of selection (39). Using this ‘sensitivity’ approach for an age-structured population (20, 37 – 38, 40 – 42), the force of selection acting on a mutant allele can be written as:

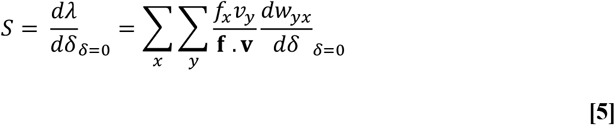

where **f** and **v** are the vector of asymptotic frequencies and the vector of inclusive reproductive values for the different age classes in the resident population. The term *f*_*x*_ denotes the asymptotic frequency of age class *x*, and **f** is the dominant right eigenvector of the demographic projection matrix (***A***). In this model, the term *v*_*x*_ represents the inclusive reproductive value of age class *x*, and is instead derived from an inclusive fitness matrix (***W***) that decomposes the demographic projection matrix into inclusive fitness contributions between age classes. Therefore, **v** is the dominant left eigenvector of ***W***. Thus, the growth rate of the mutant population, λ, represents an inclusive fitness growth rate of the allele. Finally, the term *w*_*yx*_ represents the class *y* offspring of a class *x* individual (genetic offspring equivalents). Therefore, *dw*_*yx*_ represents the difference in the contribution of an individual age *x* to individuals aged *y* in the mutant population compared to the resident population. Overall, the sign of *S* predicts the direction of selection on the mutant allele with respect to the resident population wild type allele, whilst the magnitude of *S* conveys information about the force of selection (2, 20).

### The inclusive fitness force of selection on survival

A mutant allele that alters the survival rate between age *x* and *x* + 1 changes inclusive fitness contributions between age class according to the following (see **Supplementary Information Appendix B**):

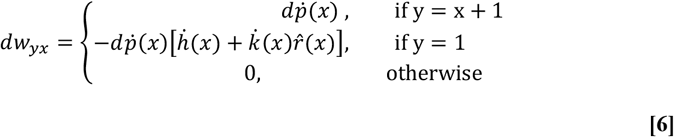

where 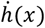 is the proportion of offspring after dispersal at the local patch that are the direct and indirect contributions of a focal individual aged 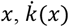 is the proportion of offspring that are born due to the genotypes of other individuals on the patch, and 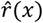 is the relatedness of an individual aged *x* to the offspring of other patch mates (see **Supplementary Information Appendix A**). As we assume mortality occurs between breeding seasons, a focal individual’s contributions to the survival and reproduction of other age classes are only affected at *x* + 1, not in the current breeding season.

Let *S*_*p*_(*x*) be the component of the force of selection due the effect of a mutant allele on the survival rate between age *x* and *x* + 1. Using equations **[5]** and **[6]**, in a stationary population with limited dispersal and social interactions between individuals, this can be written as:

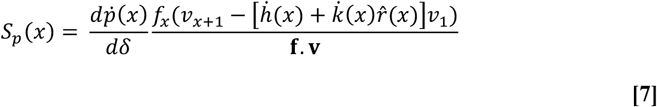

Equation **[7]** shows that the overall direction of the force of selection acting on a mutant allele that affects the survival rate between age *x* and *x* + 1 is a balance of two forces: the inclusive reproductive value at age *x* + 1 *vs* the reproductive value of offspring (displaced by the survival of the focal individual) that have varying relatedness to the focal individual aged *x*. The term **f. v** acts to scale the forces of selection in terms of generation time (2, 20).

### The inclusive fitness force of selection on reproduction

A mutant allele that alters reproduction at age *x* changes inclusive fitness contributions between age class according to the following (see **Supplementary Information Appendix B and D**):

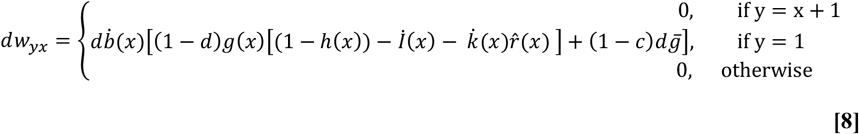

Then, let *S*_*m*_(*x*) be the component of the force of selection due the effect of a mutant allele on reproduction at age *x*. Using **[5]** and **[8]**, in a stationary population with limited dispersal and social interactions between individuals, this can be written as:

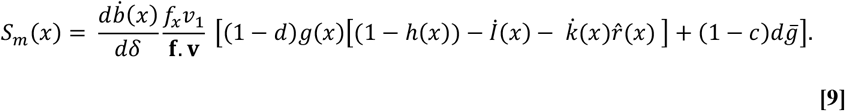

where

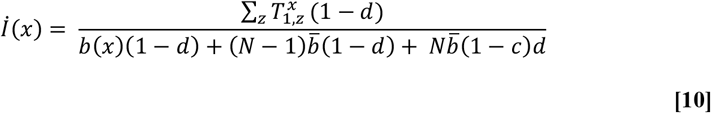

is the fraction of all offspring at the local patch after dispersal that exist due to indirect effects of the genotype of a focal individual aged *x*. Equation **[9]** shows that the overall force of selection acting on a mutant allele that affects the rate of reproduction at age *x* is also comprised of two components: *(i)* the effect of the allele on the probability of establishment of different types of offspring onto the local patch and *(ii)* the effect of the allele on the dispersing offspring that are part of the direct fitness of the focal individual aged *x*. Selection for effect *(ii)* will always be positive; however, selection for effect *(i)* will depend on the relative weights each class of offspring contributes to the overall effect. In this model, an increase in direct reproduction is, all else being equal, beneficial for the direct fitness of a focal individual, but detrimental to the indirect fitness of the focal individual.

### Applications of the model

Equations **[7]** and **[9]** provide general solutions for age-specific inclusive fitness forces of selection on individual survival and reproduction in group structured populations. To visualise the results, we consider two hypothetical populations of iteroparous individuals with social interactions (Fig 1, Fig 2). For each, we consider background demography described by age-specific vital rates, *p*(*x*) and *b*(*x*). We parameterise mortality risk at age *x* using the Siler model (43):

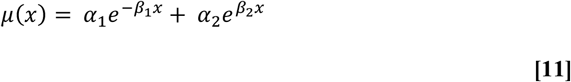

**Figure 2.**
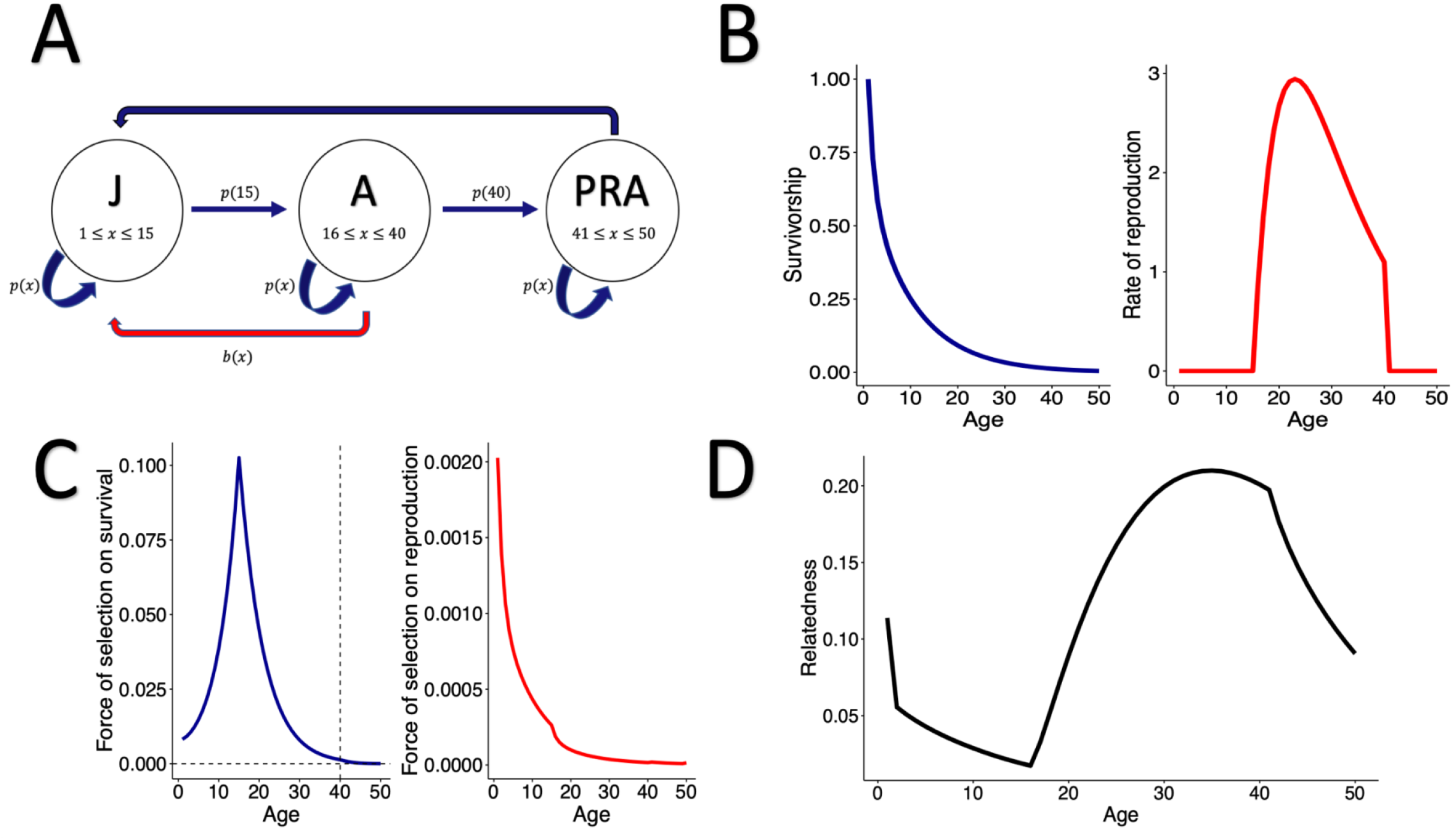
Age specific forces of selection in a social population with post-reproductive help. A) A hypothetical population of iteroparous individuals classified into three life cycle stages: juvenile (J), reproductive adult (A), and post-reproductive adult (PRA). The red arrow from A to J represents the reproduction of adult individuals, whereas the dark blue arrow from PRA to J represents the social contributions from post-reproductive adults to the survival of juveniles. B) The background vital rates of survivorship and reproduction of the model social population. Survival probability at age *x* is produced from a Siler model (**[11]**) with parameters: *α*_1_ = 0.4, *β*_1_ = 0.6, *α*_2_ = 0.1, *β*_2_ = 0 (See SOM for further details). Reproduction at age *x* is modelled according to **[13]** with parameters: *ε* = 15, *φ* = 0.125, and *κ* = 40 (SOM). C) The forces of selection acting on survival at age *x* increases during the juvenile period and then decreases but remains above zero in the post-reproductive period. The force of selection acting on reproduction at age *x* is weaker than the force of selection acting on survival and declines from birth. Other demographic parameters to produce these forces of selection were set to *c* = 0, *d* = 0.5, *N* = 4 and *ω* = 50. (see **Model** and SOM D) The relatedness of an individual aged *x* to another random individual on the patch declines throughout the juvenile (pre-reproductive) window, and then increases during adult reproduction before declining again as reproduction ceases.

**Figure 3.**
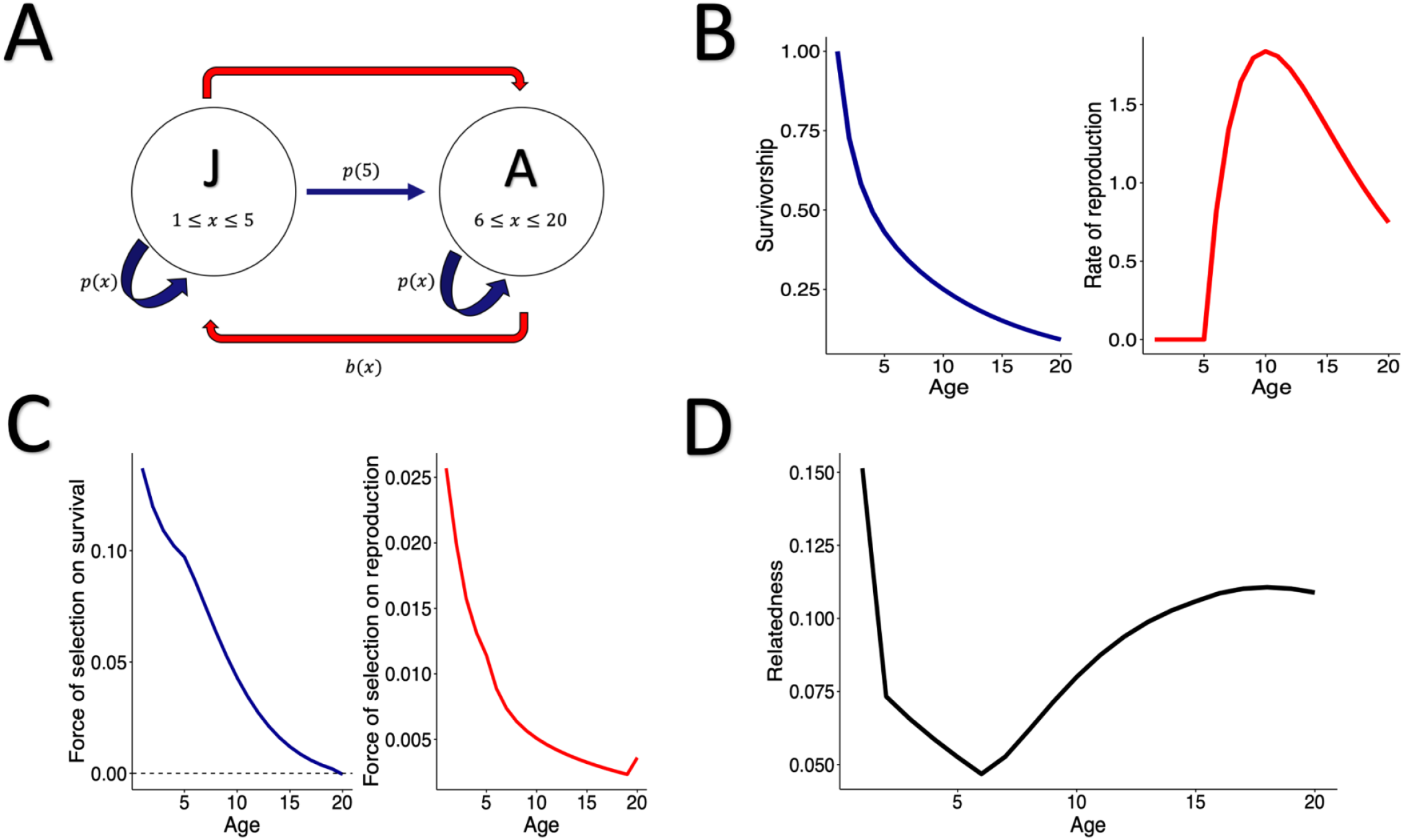
Age specific forces of selection in a social population with pre-reproductive help. A) A hypothetical population of iteroparous individuals with two lifecycle stages: juvenile (J) and reproductive adult (A). The red arrow from J to A represents the social contributions from juveniles to the reproduction of adults. Note that here, help is in the currency of reproduction, rather than survival (See Fig. 2A). B) The background vital rates of survivorship and reproduction. Survival at age *x* is produced from a Siler model (**[11]**) with parameters: *α*_1_ = 0.4, *β*_1_ = 0.6, *α*_2_ = 0.1, *β*_2_ = 0. Rate of reproduction at age *x* is modelled according to **[13]** with parameters: *ε* = 5, *φ* = 0.2, and *κ* = 21. C) The force of selection acting on survival at age *x* declines from birth. The force of selection acting on reproduction at age *x* is weaker than the force of selection on survival and also declines from birth but then increases in the final age class. Other demographic parameters to produce these forces of selection were set to *c* = 0, *d* = 0.5 and *N* = 4 and *ω* = 20. D) The relatedness of an individual aged *x* to another random individual on the patch declines throughout the juvenile period, and then increases during adult reproduction.

The probability of survival at age *x, p*(*x*), is therefore equal to *e*^−*μ*(*x*)^. The probability of survival to age *x* (*l*(*x*)) is then 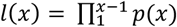, with *l*(1) = 1. As we assume all patches have no breeding positions available at the start of each breeding seasons (*i*.*e*., a density-dependent stationary population), we can calculate the asymptotic frequency (*f*_*x*_) of each age class as

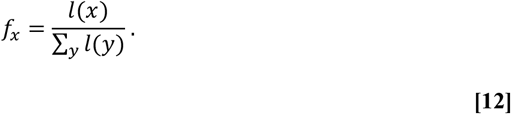

We then parameterise individual rate of reproduction at age *x* as:

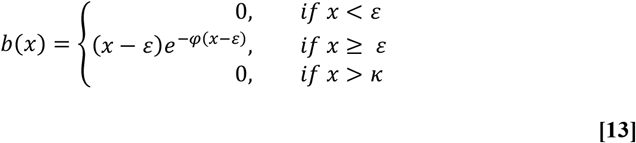

where *ε* designates the age of reproductive maturity, *κ* represents an age at which reproduction ceases, and *φ* modulates the shape of reproduction across age classes.

Fig. 1*A* and Fig. 2*A* illustrate the life cycles of the two hypothetical social populations. Fig. 1*A* considers a population with post-reproductive individuals providing care for juveniles, as seen in humans (44), orcas (45), and Asian elephants (46). Fig. 2*A* considers a population with juvenile individuals providing help to the reproduction adult breeders, as is found in many cooperatively-breeding species (47). Fig. 1*B* and Fig. 2*B* display the modelled survivorship and reproduction as a function of individual age. We then apply our methodology (see **Model and Appendix C)** to partition these vital rates into inclusive fitness contributions between age classes and compute a fitness matrix (**W**) with elements described in **[1]**. Fig. 1*C* and Fig. 2*C* show the forces of selection acting on survival and reproduction at age *x* in these hypothetical social populations according to equations **[7]** and **[9]**.

We show that the force of selection acting on survival in social populations is not necessarily constant before maturity, as predicted by classical theory (2). The exact pattern depends on whether pre-reproductive individuals gain indirect fitness through transfers or not. When juveniles do not engage in helping behaviour, the force of selection increases in the juvenile period as relatedness to newborn offspring decreases with increasing juvenile age (Fig. 1*C*; Fig.1*D*). This decline in local relatedness facilitates a more ‘selfish’ force of selection on survival throughout the juvenile period. On the other hand, when juveniles provide help to adult reproduction, the force of selection on survival generally decreased from the age at which indirect fitness was first accrued (Fig. 2*C*; Fig. S3), rather than the age of first reproduction. In both examples, the force of selection on survival then declines throughout adulthood as future inclusive reproductive value declines and the relatedness to newborn offspring increases. When post-reproductive adults continue to accrue indirect fitness, the force of selection on survival can remain above zero in post-reproductive age classes (Fig. 1*C*; Fig. S1). The magnitude of the force of selection is greater in post-reproductive age classes when juvenile dispersal is lower (and so there is higher local relatedness) and the magnitude of help provided by post-reproductive individuals is higher (Fig. S1). In general, the force of selection on survival will always have a positive component until the final age at which inclusive fitness is accrued, rather than necessarily the age of last reproduction. At this age, when future survival is no longer possible, the first term on the numerator of Equation **[7]** is zero, and so, if there is some level of local relatedness (*i*.*e*. 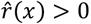), selection will favour increased mortality as it will benefit the establishment of related juveniles.

In populations with relatively long lifespans, the force of selection on reproduction was weaker than the force of selection on survival. The force of selection acting on reproduction at age *x* generally declined from birth, as predicted by Hamilton’s model (2), but not always (Fig. S4), and the decline was more rapid when the rate of dispersal was lower (Fig. S2). This more rapid decline is likely due to the greater inclusive fitness costs of increasing personal reproduction when local relatedness is higher. The force of selection on reproduction in early life is also weaker when post-reproductive adults have a more significant impact on juvenile survival. In all iterations of the model (Fig. 2*C*; Fig. S3), there was a slight increase in the force of selection acting on reproduction in the final age class, when the force of selection on rate of survival becomes negative.

## Discussion

When considering the evolution of demographic senescence, evolutionary biologists use population growth rate, *r*, as the measure of fitness (48, but see 49). The magnitude of the change in population growth rate due to an age-specific change in survival and/or reproduction generally declines with age (but see (50) for other indicators of the force of selection), and this decline facilitates the evolution of senescence (2). However, for social species, it is crucial to consider explicitly the inclusive fitness of individuals as the quantity that natural selection seeks to maximise (10). Indeed, the change in inclusive fitness due to an age-specific change in individual survival and/or reproduction must consider the combined effect on all individuals that are affected by the change (24). Here, we show that, in an age-structured model for patch-structured social populations, considering the inclusive fitness effects of an allele significantly alters the form of the forces of selection acting on age-specific survival rate and rate of reproduction.

Our framework provides several key insights into the force of selection acting on survival and reproduction in social species. First, the force of selection acting on the survival rate of that age class is the product of future inclusive reproductive value (IRV), rather than conventional reproduction value (RV (48)), and the asymptotic frequency (stationary age distribution) of that age class. Since IRV remains above zero after reproduction ceases, if post-reproductive adults continue to accrue indirect fitness benefits, selection on survival of post-reproductive age-classes does not necessarily go to zero as in Hamilton’s model (2). Importantly, this finding provides a formal inclusive fitness framework for the ‘grandmother hypothesis’ (51 – 52), supporting work that has suggested indirect fitness benefits are essential to sustained post-reproductive lifespan (23, 31). In our framework, the force of selection on survival of social species will remain non-zero until there is no future IRV. At this point, if there is some local relatedness, the force of selection on increased survival will be negative. Combined with an increase in the force of selection on reproduction at a ‘final age class’, a kin-selected terminal investment strategy, in which it pays to invest heavily in reproduction at the expense of survival to maximise the establishment of kin, may be favoured (19).

The incorporation of age-specific indirect fitness into the evolutionary theory of senescence means that selection on survival before maturity is not necessarily constant (Fig. 1*C*; Fig. 2*C*). This difference occurs because of the balance between the future IRV of the individual and the IRV of newborns displaced by increased survival. If relatedness to other individuals declines throughout the juvenile period as a focal individual ages, and the focal individuals own IRV increases as they approach maturity, the balance in Equation **[7]** is weighed more heavily towards the first term, and the force of selection on increased survival will increase. On the other hand, in populations where juveniles help and accrue indirect fitness, the force of selection on survival will declined from the age at which indirect fitness is first gained. This result implies that, in species with pre-reproductive help, senescence should start from the age at which inclusive fitness is first gained, rather than the age of first reproduction, as in conventional models (2, 20).

An inclusive fitness force of selection acting on reproduction depends on the costs and benefits associated with increasing personal reproduction. In our framework, selection for increased reproduction will always have a positive component due to the increased probability of an offspring (whether philopatric or dispersive) establishing on to a patch. However, the subsequent decrease in probability of other locally produced offspring establishing on to the patch reduces the magnitude of the force of selection acting on reproduction. This result may be especially important for groups experiencing strong competition over resources (12). For example, a negligible force of selection on reproduction may favour reproductive restraint by some individuals within cooperatively-breeding groups, when access to reproduction is limited and inclusive fitness costs of increasing personal reproduction would be substantial (32).

Our framework builds on previous work that has made significant ground in incorporating social effects into the evolutionary theory of senescence. Lee’s (23) model showed that the force of selection acting on age-specific mortality can be modified by intergenerational transfers of resources. However, kin selection did not enter the formal model as no explicit spatial structured was considered. Here, by explicitly considering a patch structured population with dispersal, we allow for variation in relatedness and thus a larger breadth of possible kin selection effects to be considered. Ronce & Promislow (20) derived analytical solutions that provided the baseline framework for the model here, showing that the force of selection on increased survival includes a negative component driven by the displacement of offspring from establishing on the local patch. This term is similar to the negative term in **[7]**; however, our framework also explicitly considers the impact of survival on the establishment of other locally produced offspring. By only considering single individuals on a patch, social interactions in Ronce & Promislow’s model were limited to kin competition between parent and offspring over residency on the patch. Here, by including multiple individuals on the patch, we can also incorporate social effects into the form of the force of selection on reproduction (**[10]**). Finally, Moorad & Nussey (53) took a quantitative genetics approach to add indirect genetic effects, explicitly considering maternal effect senescence, but modelled no explicit demography. A combination of explicit demography, as modelled here, and quantitative genetics could prove a major future step.

The framework we present here provides a base to expand our understanding of senescence across social species. For example, previous work has found mixed evidence for extended lifespan in cooperative breeders (54 – 57), and some evidence for differences in rates of senescence between cooperative and non-cooperative breeders (58). Previous theory suggests that it is longer life and overlapping generations that initially favour cooperation (26), but also that a delayed age of first reproduction as a result of queuing for reproduction might be a self-reinforcing mechanism for extended lifespan in cooperative breeders (59). However, multiple other facets of the demography of cooperative breeding systems, including the process of group formation (60), the structure of dominance hierarchies (61) and levels of reproductive skew (62) all have the potential to play a role in determining lifespan and rates of senescence. All have the potential to contribute to the shape of the age class asymptotic frequency and inclusive reproductive value distributions that, as we have shown here, underpin inclusive fitness forces of selection. Our model provides a framework to stimulate further theoretical work for how these features of cooperative breeding systems may impact the evolution of lifespan and senescence.

Here, we focused on how cooperative interactions between members of a group can alter age-specific inclusive fitness forces of selection. However, in many groups, competitive interactions over limited resources are also rife. In our model, transfers between age classes reflect the net effect of the presence of an individual in one age class on the survival and reproduction of an individual in another age class. If the net effect is negative, then the genetic offspring transfer is also negative. For example, consider again the social system illustrated in Fig. 1. Instead of post-reproductive individuals having a positive effect of the survival of juveniles, let us instead imagine a scenario in which the presence of post-reproductive individuals is harmful to the survival of juveniles. An allele that increases the rate of survival in such post-reproductive individuals will be selected against due to the inclusive fitness costs imposed from the negative effects on related juvenile individuals, potentially hastening the evolution of more rapid senescence. Finally, in our model, we only considered indirect fitness returns from social interactions. In many cooperative breeding systems, however, direct fitness returns from social interactions can be the main driver for alloparental care (47). Some form of direct fitness benefits could be incorporated into the model by delaying the age at which returns from social interactions are realised, as hypothesised by group augmentation theory (63).

In summary, recent research has focused on the potential for social interactions to drive variation in senescence across species (1, 64). The model we present here shows that when inclusive fitness consequences of increasing individual survival or reproduction are considered, age-specific forces of selection can vary markedly from previous asocial models. Our results thus support the hypothesis that sociality can shape patterns of senescence in nature. Further theoretical, empirical and comparative studies are now needed to determine the amount of variation in senescence patterns that can be explained by social modes of life.

## Acknowledgements

We thank A. Leeks, J. Park, M. Patel and T. Scott for helpful discussions and feedback. MJR was supported by funding from the Biotechnology and Biological Sciences Research Council (BBSRC) (BB/M011224/1). RSG was supported by a NERC Independent Research Fellowship (NE/M018458/1).

## Notes

### Competing Interest Statement

The authors have declared no competing interest.

