## Supplementary information for "Inclusive fitness forces of selection in an age-structured population"

Corresponding authors: Mark Roper

**This PDF file includes:**

Appendix A: Supplementary methods for relatedness calculations

Appendix B: Supplementary methods for analytical solutions in the main text

Appendix C: Supplementary methods for numerical applications and R code for numerical solutions

Appendix D: Further derivations for ${dw}_{1x}$

Figures S1-4

References for Supplementary Information

**Other supplementary materials for this manuscript include the following:**

R Code for producing figures (available on request)

**Appendix A: Supplementary methods for relatedness calculations**

In order to quantify indirect genetic contributions, it is essential to consider the relatedness between different age classes of individuals in the population. The relatedness of a focal individual aged $x$ to other individuals on the patch, including themselves, can then be described as:

$$r\left( x \right)= \frac{1}{N}+ \frac{N-1}{N}\hat{r}\left( x \right) .$$

**[A1]**

Then, let $r_{yx}$ denote the probability that an allele sampled randomly from a given locus in an individual aged $x$ is identical by descent (IBD) to an allele sampled randomly from the same locus in an individual aged $y$ (1 – 6). The term $\hat{r}\left( x \right)$ represents the average relatedness of a breeding individual aged $x$ to another random breeder on the same patch (1 – 2), which is equivalent to the mean relatedness of a focal individual aged $x$ across all age classes ($\hat{r}\left( x \right)=\bar{r_{yx}}$). Given the assumption of haploid genetics and asexuality, $\hat{r}\left( x \right)$is therefore also the relatedness of a focal individual aged $x$ to the offspring of the other individuals on the patch. Under the assumption of infinite patches, any immigrants arriving at the focal patch will not have any relatives when they arrive, and the relatedness of individuals on the patch of any age to these immigrants is equal to 0.

Let us define $h\left( x \right)$ as the proportion of offspring after dispersal at the local patch that are the offspring (not partitioned into inclusive fitness contributions) of a focal individual aged $x$:

$$h\left( x \right)= \frac{b(x)(1-d)}{b\left( x \right)(1-d)+(N-1)\bar{b}(1-d)+N\bar{b}d(1-c)}$$

**[A2]**

where $\bar{b}$ represents the average rate of reproduction. For simplicity, we assume no demographic stochasticity within patches (see **Discussion**). Then, let $k\left( x \right)$ define the proportion of offspring after dispersal at the local patch that are the demographic offspring of other individuals on the patch besides the focal individual aged $x$:

$$k\left( x \right)= \frac{(N-1)\bar{b}(1-d)}{b\left( x \right)1-d)+(N-1)\bar{b}(1-d)+N\bar{b}d(1-c)}$$

**[A3]**

Using equations **[A2]** and **[A3]**, we can describe the relatedness between an individual aged $x$ to a different individual on the patch aged $y$ as a function of both individual’s ages:

$$r_{yx}=\left\{ \begin{aligned} h\left( x-y \right)+k\left( x-y \right)\hat{r}(x-y), &y<x \\ \left( 1-d \right)^{2}\left[ \bar{h}^{2} +(1-\bar{h}^{2})\hat{r}(1) \right] , &y=x \\ h\left( y-x \right)+k(y-x)\hat{r}(1), &y>x \end{aligned} \right. .$$

**[A4]**

First, consider the case when the individual of age $x$ is older than the individual of age $y$ (top row of **[A4]**). The individual aged $y$ was born $x-y$ breeding seasons ago, when the individual aged $x$ was $x-y$ years old. At age $x-y$, the proportion of offspring at the local patch after dispersal that are the offspring of an individual aged $x-y$ is defined as $h\left( x-y \right)$. Therefore, with probability $h\left( x-y \right)$, the individual aged $y$ is the offspring of the individual aged $x$ from $x-y$ breeding seasons ago, and thus the relatedness between the two individuals is one. Then, let $k\left( x-y \right)$ define the proportion of offspring at the local patch after dispersal$x-y$ breeding seasons ago that were the offspring of other individuals on the patch. With probability $k\left( x-y \right),$therefore, the individual aged aged $y$ was born to another individual on the patch. Therefore, the relatedness of the individual aged $x$ to the individual aged $y$ is equal to the relatedness of an individual aged $x-y$ to a random offspring born locally to the patch, which is equal to the relatedness of an individual aged $x-y$ to another random individual on the patch ($\hat{r}(x-y)$). The remaining proportion of offspring at the patch after dispersal $x-y$ breeding seasons ago ($1-h\left( x-y \right)-k(x-y)$) were from elsewhere in the population and thus relatedness is 0.

Second, consider the case when both individuals are the same age (second row of **[A4]**). The probability that both are local to the patch is $\left( 1-d \right)^{2}$. If both individuals are born locally, we then have to consider the probability that both individuals were born to the same mother, and thus are siblings related by 1. If the average proportion across age classes of offspring that are born to an individual is $\bar{h}$, then the probability that two offspring born $x$ breeding seasons ago were born to the same mother is equal to $\bar{h}^{2}$. One minus $\bar{h}^{2}$is then the probability that these two locally born offspring $x$ breeding seasons ago were born to different mothers, in which case the relatedness of an individual aged $x$ to a same aged individual is equal to the relatedness of an individual to a random member of the patch at age 1 when the focal individual established onto the patch ($\hat{r}(1)$). The final scenario (bottom row of **[A4]**) considers the case when the individual aged $y$ is older than the individual aged $x$. In this case the logic is the opposite to the case when the individual aged $x$ is older than the individual aged $y$.

To calculate $\hat{r}\left( x \right)$, the average relatedness of an individual aged $x$ to another individual on the patch, we need to calculate the average relatedness of individuals aged $x$ to all other age classes. Using each possible relatedness between age classes (**[A4]**), we can do this by weighting each age class specific relatedness term by the asymptotic frequencies of the relevant age classes:

$$\hat{r}(x)=\left( \sum^{y<x} f_{y}\left[ \left( h(x-y \right)+k(x-y)\hat{r}(x-y) \right] \right)+ f_{x}\left( 1-d \right)^{2}\left( \bar{h}^{2}+\left( 1-\bar{h}^{2} \right)\hat{r}(1) \right)+ \left( \sum_{y=x+1}^{y=\omega} f_{y}\left[ h(y-x)+k(y-x)\hat{r}(1) \right] \right)$$

**[A5]**

**Deriving** $\hat{\boldsymbol{r}}\left( \boldsymbol{1} \right)$

To find a general solution for $\hat{r}\left( 1 \right)$, which is the relatedness of an individual aged 1 to another random breeder on the patch, let us consider a case of a population with 3 age classes ($\omega=3)$. Using the logic that $x=1$ is the first age class and therefore $y$ cannot be younger than $x$, $\hat{r}\left( 1 \right)$ with 3 age classes becomes:

$$\hat{r}\left( 1 \right)=f_{1}\left( 1-d \right)^{2}\left[ \bar{h}^{2}+ \hat{r}\left( 1 \right)\left( 1-\bar{h}^{2} \right) \right]+ \sum_{y=2}^{3} f_{y}\left[ h\left( y-x \right)+k\left( y-x \right) \right]\hat{r}\left( 1 \right)$$

**[A6]**

Expanding the summation term, this becomes:

$$\hat{r}\left( 1 \right)=f_{1}\left( 1-d \right)^{2}\left[ \bar{h}^{2}+ \hat{r}\left( 1 \right)\left( 1-\bar{h}^{2} \right) \right]+ f_{2}\left[ h\left( 1 \right)+k\left( 1 \right)\hat{r}\left( 1 \right) \right]+ f_{3}\left[ h\left( 2 \right)+k\left( 2 \right)\hat{r}\left( 1 \right) \right]$$

**[A7]**

Expanding out each term, this becomes:

$$\hat{r}\left( 1 \right)= f_{1}\left( 1-d \right)^{2}\bar{h}^{2}+ f_{1}\left( 1-d \right)^{2}\hat{r}\left( 1 \right)\left( 1-\bar{h}^{2} \right)+ f_{2} h\left( 1 \right)+f_{2}k\left( 1 \right)\hat{r}\left( 1 \right)+f_{3} h\left( 2 \right)+ f_{3}k(2)\hat{r}\left( 1 \right)$$

**[A8]**

### Factoring on the RHS by $\hat{r}\left( 1 \right)$, this becomes:

$$\hat{r}\left( 1 \right)= \hat{r}\left( 1 \right)\left[ f_{1}\left( 1-d \right)^{2}\left( 1-\bar{h}^{2} \right)+ f_{2}k\left( 1 \right)+f_{3}k(2) \right]+f_{1}\left( 1-d \right)^{2}\bar{h}^{2}+ f_{2} h\left( 1 \right)+f_{3} h\left( 2 \right)$$

**[A9]**

Re-arranging, and factoring on the LHS by $\hat{r}\left( 1 \right)$ this becomes:

$$\hat{r}\left( 1 \right)\left[ 1-\left[ f_{1}\left( 1-d \right)^{2}\left( 1-\bar{h}^{2} \right)+ f_{2}k\left( 1 \right)+f_{3}k(2) \right] \right]= f_{1}\left( 1-d \right)^{2}\bar{h}^{2}+ f_{2} h\left( 1 \right)+f_{3} h\left( 2 \right)$$

**[A10]**

Dividing both sides by $\left[ 1-\left[ f_{1}\left( 1-d \right)^{2}\left( 1-\bar{h}^{2} \right)+ f_{2}k\left( 1 \right)+f_{3}k(2) \right] \right]$, this becomes:

$$\hat{r}\left( 1 \right)= \frac{f_{1}\left( 1-d \right)^{2}\bar{h}^{2}+ f_{2} h\left( 1 \right)+f_{3} h\left( 2 \right)}{1- \left[ f_{1}\left( 1-d \right)^{2}\left( 1-\bar{h}^{2} \right)+ f_{2}k\left( 1 \right)+f_{3}k(2) \right]}$$

**[A11]**

Finally, to generalise for all possible number of age classes, we can re-write **[A11]** as

$$\hat{r}\left( 1 \right)= \frac{f_{1}\left( 1-d \right)^{2}\bar{h}^{2}+ \sum_{y=2}^{\omega} f_{y}h(y-1)}{1- \left[ f_{1}\left( 1-d \right)^{2}\left( 1-\bar{h}^{2} \right)+\sum_{y=2}^{\omega} f_{y}k(y-1) \right]}$$

**[A12]**

Once we have $\hat{r}\left( 1 \right)$, $\hat{r}\left( x \right)$ for all other age classes can be solved recursively.

**Appendix B: Supplementary methods for analytical solutions in the main text**

**The effect of a mutant allele that alters age-specific survival in a social population**

Let us first consider how, in a resident population with limited dispersal and social interactions, a mutant allele that affects survival at age $x$ will alter the number of class-$y$ offspring of a focal individual aged $x$. First, the most obvious effect of this allele is to change the individual’s probability of survival to the next breeding season, which is$d\dot{p}(x)$. A change in survival will also alter the contributions a focal individual aged $x$ makes to the offspring class, $w_{1x}$. For example, if the mutant allele increases survival at age $x$, then there is a greater chance the focal individual survives to age $x+1$, and this subsequently reduces the probability that an offspring at the focal patch after dispersal will establish onto the patch before the next breeding season. Four classes of offspring will exist at the focal patch after dispersal: 1) the offspring of a focal individual aged $x$, 2) the offspring of other individuals on the patch that exist due to the genotype of a focal individual aged $x$, 3) the offspring of other individuals on the patch that don’t owe their existence to the genotype of a focal individual aged $x$, and 4) offspring from elsewhere in the population. As we are interested in the inclusive fitness effect of the mutant allele, we must consider the fates of all the offspring that are impacted by the effect of the allele (7).

We can consider the first two sets of offspring together and ask how a change in survival at age $x$ alters the direct and indirect production of offspring of a focal age $x$ individual (working showed below).

$$\frac{{dw}_{1x}(1,2)}{d\dot{p}(x)}=\dot{F}\left( x \right)\left[ \left( 1-d \right)g\left( x \right)+\left( 1-c \right)d\bar{g} \right]- \dot{F}\left( x \right)\left[ \left( 1-d \right)g^{'}(x)+\left( 1-c \right)d\bar{g} \right]$$

**[B1]**

with $g^{'}(x)$ displaying that the effect of the allele is to alter the probability that the direct and indirect offspring of the individual aged $x$ establish on to the patch. **[B1]** can be worked through and simplified as:

$$\frac{{dw}_{1x}(1,2)}{d\dot{p}(x)}=\dot{F}\left( x \right)\left( 1-d \right)g\left( x \right)+\dot{F}\left( x \right)\left( 1-c \right)d\bar{g}-\dot{F}\left( x \right)\left( 1-d \right)g^{'}\left( x \right)-\dot{F}\left( x \right)\left( 1-c \right)d\bar{g}$$

$$=\dot{F}\left( x \right)\left( 1-d \right)g\left( x \right)-\dot{F}\left( x \right)\left( 1-d \right)g'(x)$$

$$=\dot{F}(x)(1-d)\left[ g\left( x \right)-g'(x) \right]$$

$$=\dot{F}(x)(1-d)\left[ \frac{1-p\left( x \right)+(N-1)(1-\bar{p})}{b\left( x \right)\left( 1-d \right)+\left( N-1 \right)\bar{b}\left( 1-d \right)+ N\bar{b}\left( 1-c \right)d}- \frac{1-p^{'}\left( x \right)+(N-1)(1-\bar{p})}{b\left( x \right)\left( 1-d \right)+\left( N-1 \right)\bar{b}\left( 1-d \right)+ N\bar{b}\left( 1-c \right)d} \right]$$

$$=\dot{F}(x)(1-d)\left[ \frac{-d\dot{p}(x)}{b\left( x \right)\left( 1-d \right)+\left( N-1 \right)\bar{b}\left( 1-d \right)+ N\bar{b}\left( 1-c \right)d} \right]$$

$$= -d\dot{p}(x)\left[ \frac{\dot{F}(x)(1-d)}{b\left( x \right)\left( 1-d \right)+\left( N-1 \right)\bar{b}\left( 1-d \right)+ N\bar{b}\left( 1-c \right)d} \right]$$

Finally, let $\dot{h}\left( x \right)=\frac{\dot{F}(x)(1-d)}{b\left( x \right)\left( 1-d \right)+\left( N-1 \right)\bar{b}\left( 1-d \right)+ N\bar{b}\left( 1-c \right)d}$be defined as the proportion of offspring at the focal patch after dispersal that are born due the genotype of a focal individual aged $x$. Note, $\dot{h}\left( x \right)$ is different from $h\left( x \right)$ (see **Supplementary Information Appendix A**), as $h\left( x \right)$ does not partition the offspring with respect to inclusive fitness contributions. The relatedness of the indirect offspring has already been discounted in the calculation of $\dot{F}(x)$, and the relatedness of a focal individual to its own offspring is 1, so we can re-write **[B1]** as

$$\frac{{dw}_{1x}(1,2)}{dp(x)}=-d\dot{p}\left( x \right)\dot{h}(x)$$

**[B2]**

Let us now consider the third set of offspring and ask how a change in survival of a focal individual at age $x$ impacts the offspring of other individuals on the patch that don’t owe their existence to the genotype of a focal individual aged $x$. In the resident population, this contribution is 0. However, an increase in survival of an individual aged $x$, for example, will reduce the likelihood that any of these offspring that do not disperse will establish onto the patch before the next breeding season. We can write the average number of offspring of all other individuals on the patch, in the presence of a focal individual aged $x$, that will establish onto the local patch as

$$\left( N-1 \right)\bar{F}(1-d)g(x)$$

**[B3]**

The effect of a mutant allele that alters the survival of a focal individual aged $x$ on this expected number of offspring can then be written as

$$\frac{{dw}_{1x}(3)}{d\dot{p}(x)}= \left( N-1 \right)\bar{F}\left( 1-d \right)g\left( x \right)- \left( N-1 \right)\bar{F}\left( 1-d \right)g'\left( x \right)$$

**[B4]**

**[B4]** can then be worked through and simplified as

$$\frac{{dw}_{1x}\left( 3 \right)}{d\dot{p}\left( x \right)}= \left( N-1 \right)\bar{F}\left( 1-d \right)\left[ g\left( x \right)-g^{'}\left( x \right) \right]$$

$$=\left( N-1 \right)\bar{F}\left( 1-d \right)\left[ \frac{1-p\left( x \right)+(N-1)(1-\bar{p})}{b\left( x \right)\left( 1-d \right)+\left( N-1 \right)\bar{b}\left( 1-d \right)+ N\bar{b}\left( 1-c \right)d}- \frac{1-p^{'}\left( x \right)+(N-1)(1-\bar{p})}{b\left( x \right)\left( 1-d \right)+\left( N-1 \right)\bar{b}\left( 1-d \right)+ N\bar{b}\left( 1-c \right)d} \right]$$

$$=\left( N-1 \right)\bar{F}\left( 1-d \right)\left[ \frac{-d\dot{p}(x)}{b\left( x \right)\left( 1-d \right)+\left( N-1 \right)\bar{b}\left( 1-d \right)+ N\bar{b}\left( 1-c \right)d} \right]$$

$$= -d\dot{p}(x)\left[ \frac{\left( N-1 \right)\bar{F}\left( 1-d \right)}{b\left( x \right)\left( 1-d \right)+\left( N-1 \right)\bar{b}\left( 1-d \right)+ N\bar{b}\left( 1-c \right)d} \right]$$

Similar to the logic above, let $\dot{k}\left( x \right)=\frac{\left( N-1 \right)\bar{F}\left( 1-d \right)}{b\left( x \right)\left( 1-d \right)+\left( N-1 \right)\bar{b}\left( 1-d \right)+ N\bar{b}\left( 1-c \right)d}$ be defined as the proportion of offspring at the focal patch after dispersal that are average direct and indirect offspring of all other individuals bar the focal individual aged $x$. These offspring are related to the focal individual by $\hat{r}(x)$ and so the above becomes

$$\frac{{dw}_{1x}\left( 3 \right)}{d\dot{p}\left( x \right)}= -d\dot{p}(x)\dot{k}(x)\hat{r}(x)$$

**[B5]**

Given our assumptions of an infinite population, we can assume that relatedness of any individual on a patch to offspring that have dispersed from elsewhere will be equal to zero. Therefore, the relatedness of a focal individual aged $x$ to the proportion of offspring after dispersal that were not born locally on the patch is zero. Thus, there is an overall balance of the effect of the mutant allele on a focal individual of age $x$’s production of newborns weighted on one side by locally produced offspring (with varying relatedness) and on the other side by dispersed offspring. The total effect of a mutant allele that alters age-specific survival on the production of offspring can then be summed as

$$\frac{{dw}_{1x}}{d\dot{p}\left( x \right)}=-d\dot{p}\left( x \right)\dot{h}\left( x \right) -d\dot{p}\left( x \right)\dot{k}\left( x \right)r_{1x}= -d\dot{p}\left( x \right)[\dot{h}\left( x \right)+\dot{k}\left( x \right)\hat{r}(x) ]$$

**[B6]**

The overall effect ${(dw}_{yx}$ for all y) of a mutant allele that alters age-specific survival is then shown in **[6]** in the main text.

**The effect of a mutant allele that alters age-specific reproduction in a social population**

Let us now consider how a mutant allele that affects reproduction at age $x$ will alter the class-$y$ offspring a focal individual aged $x$ in our social population. First, we assume for simplicity that a change in reproduction of a focal individual aged $x$does not alter the individual’s probability of survival to the next breeding season, or its contributions to the survival of other individuals alive on the patch. These are obvious extensions for future iterations of the model (**see Discussion in main text)**. We therefore limit the effects of a change in reproduction to altering the contributions a focal individual aged $x$ makes to the offspring class, $w_{1x}$. There are four different types of offspring to consider: 1) the offspring of a focal individual aged $x$ that exist due to its own genotype, 2) the offspring of other individuals on the patch that exist due to the genotype of a focal individual aged $x$, 3) the offspring of other individuals on the patch that don’t owe their existence to the genotype of a focal individual aged $x$, and 4) offspring from elsewhere in the population. Again, as we are interested in the inclusive fitness effect of the mutant allele, we must consider the fates of all the offspring that are impacted by the effect of the allele (7).

The inclusive fitness effects of a mutant allele that causes a change in the direct rate of reproduction of a focal individual aged $x$ for each class of offspring can be displayed as follows:

$$\frac{{dw}_{1x}(1)}{d\dot{b}(x)}=\dot{b}\left( x \right)\left[ \left( 1-d \right)g\left( x \right)+\left( 1-c \right)d\bar{g} \right]- \dot{b}'\left( x \right)\left[ \left( 1-d \right)g^{'}(x)+\left( 1-c \right)d\bar{g} \right]$$

**[B7]**

$$\frac{{dw}_{1x}(2)}{d\dot{b}(x)}=\sum_{z} T_{1,z}^{x}\left( 1-d \right)g\left( x \right)- \sum_{z} T_{1,z}^{x}\left( 1-d \right)g'\left( x \right)$$

**[B8]**

$$\frac{{dw}_{1x}(3)}{d\dot{b}(x)}=\left( N-1 \right)\bar{F}\left( 1-d \right)g\left( x \right)- \left( N-1 \right)\bar{F}\left( 1-d \right)g'\left( x \right)$$

**[B9]**

with prime notation displaying that the explicit effects of the allele. Above, **[B7]** considers the effect of the allele on the focal individual’s direct production of offspring, **[B8]** the effect of the allele on the indirect offspring of focal, and **[B9]** the effect on offspring born to other individuals on the patch not due to the genotype of focal, but whom focal might be related to more than the population average (zero). Again, individuals that disperse from elsewhere in the population to the focal patch are assumed to be related to any individual on the patch by zero, and so the inclusive fitness effect of the allele with regards to the fourth class of offspring is also equal to zero. Furthermore, given our assumption of infinite patches, the effect of the allele on the second and third classes of offspring is limited to those offspring which do not disperse *i.e.* compete for a site at the local patch. The simplification of **[B7 – B9]** follows the same logic as **[B1 – B5]**. The resulting derivations are lengthy and so are available in the separate appendix **D**. The overall effect of the mutant allele that causes a change in the rate of reproduction of a focal individual aged $x$ is the sum of the effects **[B7 – B9]** and can be expressed as:

$$\frac{{dw}_{1x}}{d\dot{b}(x)}= d\dot{b}(x)\left[ \left( 1-d \right)g\left( x \right)\left[ (1-h\left( x \right))-\dot{I}\left( x \right)- \dot{k}(x)\hat{r}(x) \right]+\left( 1-c \right)d\bar{g} \right]$$

**[B10]**

The overall effect ${(dw}_{yx}$ for all y) of a mutant allele that alters age-specific reproduction is then shown in **[8]** in the main text.

**Appendix C: Supplementary methods for numerical applications and R code**

**Numerical application methods**

To compute the inclusive fitness matrix (**W)**, we first model the propensity of each age class to contribute to the survival and reproduction of each age class, including its own. These genetic offspring transfer propensities can be conceptualised as two different matrices, one for survival and one for reproduction. The entry $\mathbf{T}_{\boldsymbol{yx}}$ represents the relative propensity for age class $x$ to contribute to the survival or reproduction of age class $y.$ This propensity takes the functional form of

$$\mathbf{T}_{\boldsymbol{yx}}=T(x, y, C_{x}, C_{y})$$

**[C1]**

Where $C$ represents the stage of each age class, which could be, for example, pre-reproductive (juveniles), mature reproductive, mature non-reproductive, or post-reproductive. The form of **[C1]** allows for the relative propensity to be a function of the age and stage of both actor and recipient. Here, for simplicity, for our population with post-reproductive survival (Fig. 1) we input a 1 into each element of the **T** matrix for survival in which the column represents a post-reproductive age class and the row represents a juvenile age class. This means that every post-reproductive age class has the same relative propensity to contribute to the survival of each juvenile age class. Every other element in **T** (survival) is set to 0, and every element in **T** (reproduction) is 0. On the contrary, for our population with pre-reproductive help, we input a 1 into each element of the **T** matrix for reproduction with a column representing a pre-reproductive age class and a column representing a reproductive age class. This means every pre-reproductive age class has the same relative propensity to contribute to the reproduction of each reproductive age class. Every other element in **T** (reproduction) = 0, and every element in **T** (survival) = 0.

The next step is to model the proportions of age-specific survival and reproduction that are due to the social environment. The total social transfers received by age class $x$is $\sum_{z} T_{x+1,x}^{z}$ for survival at age $x$ and $\sum_{z} T_{1,x}^{z}$ for reproduction at age class $x$, which are modelled as proportions of the background demographic rates $p(x)$ and $b(x)$. These proportions may vary by age and stage, and different age classes within the same stage could have different proportions stripped from their survival and reproduction. For example, very young juveniles might owe their survival more to other members of the group than older juveniles. Here, we model cases were 10, 20 and 30% of juvenile survival (Fig.1; Fig. S1; Fig. S2) or 10, 20 and 30% of adult reproduction is due to the social environment. These contributions are independent of age within the stage classification *i.e.* the same proportion is stripped from all juvenile or adult stage classes within the same iteration of the model.

We can then quantify indirect fitness contributions by distributing the stripped genetic offspring equivalents that are the result of the social environment. Specifically, a focal individual aged $x$’s genetic offspring contribution to the survival of age class $z$ is:

$$T_{z+1, z}^{x}={\left( N-1 \right)f_{z}t}_{z+1,z}^{x}\hat{r}(x)$$

**[C2]**

where $(N-1)f_{z}$ is the expected number of individuals alive on the patch aged $z$ with the focal individual aged $x$, $\hat{r}(x)$ is the relatedness of a focal individual aged $x$ to a random breeder on the patch, and $t_{z+1,z}^{x}$ represents the fraction of the total social contributions to the survival of age class $z$ that are due to age class $x$, and can be written more explicitly as

$$t_{z+1,z}^{x}=\sum_{j} T_{z+1,z}^{j}\frac{f_{x}\mathbf{T}_{zx}}{\sum_{j} f_{j}\mathbf{T}_{zj}}$$

**[C3]**

The same logic applies to reproductive contributions. For each age class, these indirect genetic contributions are compiled into the inclusive fitness matrix **W**, along with the direct components of survival ($\dot{p}(x)$) and reproduction ($\dot{b}(x)$). The dominant left eigenvector of **W** is equal to the vector of inclusive reproductive value for each age class. The inclusive fitness forces of selection acting on a mutant allele that alters the rate of survival between age $x$ and $x+1$ or rate of reproduction at age $x$ can then be calculated according to **[7]** and **[9]** in the main text. Code for the numerical solutions explored in the main text is available in a separate file.

**Appendix D: Further derivations for** $\boldsymbol{dw}_{\boldsymbol{1}\boldsymbol{x}}$

The inclusive fitness effect of a mutant allele that causes a change in the direct rate of reproduction of a focal individual aged age *x* on it’s direct fitness can be displayed as

$$\frac{{dw}_{1x}(1)}{d\dot{b}(x)}=\dot{b}\left( x \right)\left[ \left( 1-d \right)g\left( x \right)+\left( 1-c \right)d\bar{g} \right]- \dot{b}'\left( x \right)\left[ \left( 1-d \right)g^{'}(x)+\left( 1-c \right)d\bar{g} \right]$$

**[D1]**

The following working displays the simplification of **[D1]:**

$$\frac{{dw}_{1x}(1)}{d\dot{b}(x)}= \dot{b}\left( x \right)\left( 1-d \right)g\left( x \right)+ \dot{b}\left( x \right)\left( 1-c \right)d\bar{g}- \dot{b}'\left( x \right)\left( 1-d \right)g^{'}(x) -\dot{b}'\left( x \right)\left( 1-c \right)d\bar{g} ,$$

$$\frac{{dw}_{1x}(1)}{d\dot{b}(x)}= \dot{b}\left( x \right)\left( 1-d \right)g\left( x \right)- \dot{b}'\left( x \right)\left( 1-d \right)g^{'}(x) +\left( 1-c \right)d\bar{g}\left[ \dot{b}\left( x \right)-\dot{b}'\left( x \right) \right],$$

$$\frac{{dw}_{1x}(1)}{d\dot{b}(x)}= \left( 1-d \right)\left[ \frac{\dot{b}\left( x \right)\left[ (1-p\left( x \right)+(N-1)\bar{p} \right]}{b\left( x \right)\left( 1-d \right)+(N-1)\bar{b}(1-d)+ N\bar{b}\left( 1-c \right)d}- \frac{\dot{b}'(x)\left[ (1-p\left( x \right)+(N-1)\bar{p} \right]}{b'(x)(1-d)+ (N-1)\bar{b}(1-d)+ N\bar{b}\left( 1-c \right)d} \right]+ \left( 1-c \right)d\bar{g}\left[ b\left( x \right)-b'(x) \right]$$

The terms inside the brackets are manipulated using algebraic subtraction and, after simplification, results in:

$$\frac{{dw}_{1x}(1)}{d\dot{b}(x)}= \left( 1-d \right)\left[ \frac{\left[ b\left( x \right)-b'(x) \right] \left[ (1-p\left( x \right)+(N-1)\bar{p} \right]\left[ (N-1)\bar{b}(1-d)+ N\bar{b}\left( 1-c \right)d \right]}{\left[ b\left( x \right)\left( 1-d \right)+(N-1)\bar{b}(1-d)+ N\bar{b}\left( 1-c \right)d \right]\left[ b'(x)(1-d)+ (N-1)\bar{b}(1-d)+ N\bar{b}\left( 1-c \right)d \right]} \right]+ \left( 1-c \right)d\bar{g}\left[ b\left( x \right)-b'(x) \right]$$

Further simplification results in the expression:

$$\frac{{dw}_{1x}(1)}{d\dot{b}(x)}= \left( 1-d \right)\left[ b\left( x \right)-b'(x) \right]\left[ \frac{\left[ (1-p\left( x \right)+(N-1)\bar{p} \right]\left[ (N-1)\bar{b}(1-d)+ N\bar{b}\left( 1-c \right)d \right]}{\left[ b\left( x \right)\left( 1-d \right)+(N-1)\bar{b}(1-d)+ N\bar{b}\left( 1-c \right)d \right]\left[ b'(x)(1-d)+ (N-1)\bar{b}(1-d)+ N\bar{b}\left( 1-c \right)d \right]} \right]+ \left( 1-c \right)d\bar{g}\left[ b\left( x \right)-b'(x) \right]$$

which using definitions in **Appendix B** is equal to:

$$\frac{{dw}_{1x}(1)}{d\dot{b}(x)}= \left( 1-d \right)\left[ b\left( x \right)-b^{'}\left( x \right) \right]\left[ g(x)(1-h(x) \right] + \left( 1-c \right)d\bar{g}\left[ b\left( x \right)-b'(x) \right]$$

which can finally be simplified to:

$$\frac{{dw}_{1x}(1)}{d\dot{b}(x)}= d\dot{b}(x)\left[ \left( 1-d \right)g(x)(1-h\left( x \right)) + \left( 1-c \right)d\bar{g} \right]$$

**[D2]**

The inclusive fitness effect of a mutant allele that causes a change in the direct rate of reproduction of a focal individual aged age *x* on the fates of offspring it indirectly produced through the reproduction of others can be displayed as

$$\frac{{dw}_{1x}(2)}{d\dot{b}(x)}=\sum_{z} T_{1,z}^{x}\left( 1-d \right)g\left( x \right)- \sum_{z} T_{1,z}^{x}\left( 1-d \right)g'\left( x \right)$$

**[D3]**

The following working displays the simplification of **[D3]:**

$$\frac{{dw}_{1x}(2)}{d\dot{b}(x)}= \sum_{z} T_{1,z}^{x}\left( 1-d \right)\left[ \frac{\left[ (1-p\left( x \right)+(N-1)\bar{p} \right]}{b\left( x \right)\left( 1-d \right)+(N-1)\bar{b}(1-d)+ N\bar{b}\left( 1-c \right)d}- \frac{\left[ (1-p\left( x \right)+(N-1)\bar{p} \right]}{b'(x)(1-d)+ (N-1)\bar{b}(1-d)+ N\bar{b}\left( 1-c \right)d} \right]$$

The terms inside the brackets are manipulated using algebraic subtraction and, after simplification, results in:

$$\frac{{dw}_{1x}(2)}{d\dot{b}(x)}= \sum_{z} T_{1,z}^{x}\left( 1-d \right)\left[ \frac{\left[ b\left( x \right)-b'(x) \right] \left[ (1-p\left( x \right)+(N-1)\bar{p} \right](1-d)}{\left[ b\left( x \right)\left( 1-d \right)+(N-1)\bar{b}(1-d)+ N\bar{b}\left( 1-c \right)d \right]\left[ b'(x)(1-d)+ (N-1)\bar{b}(1-d)+ N\bar{b}\left( 1-c \right)d \right]} \right]$$

Further simplification and using definitions in **Appendix B** results in the expression:

$$\frac{{dw}_{1x}(2)}{d\dot{b}(x)}= d\dot{b}(x)\left( 1-d \right)\left[ \frac{\left[ (1-p\left( x \right)+(N-1)\bar{p} \right]\left[ \sum_{z} T_{1,z}^{x}(1-d) \right]}{\left[ b\left( x \right)\left( 1-d \right)+(N-1)\bar{b}(1-d)+ N\bar{b}\left( 1-c \right)d \right]\left[ b'(x)(1-d)+ (N-1)\bar{b}(1-d)+ N\bar{b}\left( 1-c \right)d \right]} \right]$$

which can finally be simplified to:

$$\frac{{dw}_{1x}(2)}{d\dot{b}(x)}=- d\dot{b}(x)\left[ \left( 1-d \right)g(x)\dot{I}(x) \right]$$

**[4]**

Finally, the inclusive fitness effect of a mutant allele that causes a change in the direct rate of reproduction of a focal individual aged age *x* on the fates of other offspring born on the patch can be written as

$$\frac{{dw}_{1x}(3)}{d\dot{b}(x)}=\left( N-1 \right)\bar{F}\left( 1-d \right)g\left( x \right)- \left( N-1 \right)\bar{F}\left( 1-d \right)g'\left( x \right)$$

**[D5]**

The following working displays the simplification of **[D5]:**

$$\frac{{dw}_{1x}(3)}{d\dot{b}(x)}= \left( N-1 \right)\bar{F}\left( 1-d \right)\left[ \frac{\left[ (1-p\left( x \right)+(N-1)\bar{p} \right]}{b\left( x \right)\left( 1-d \right)+(N-1)\bar{b}(1-d)+ N\bar{b}\left( 1-c \right)d}- \frac{\left[ (1-p\left( x \right)+(N-1)\bar{p} \right]}{b'(x)(1-d)+ (N-1)\bar{b}(1-d)+ N\bar{b}\left( 1-c \right)d} \right]$$

The terms inside the brackets are manipulated using algebraic subtraction and, after simplification, results in:

$$\frac{{dw}_{1x}(3)}{d\dot{b}(3)}= \left( N-1 \right)\bar{F}\left( 1-d \right)\left[ \frac{\left[ b\left( x \right)-b'(x) \right] \left[ (1-p\left( x \right)+(N-1)\bar{p} \right](1-d)}{\left[ b\left( x \right)\left( 1-d \right)+(N-1)\bar{b}(1-d)+ N\bar{b}\left( 1-c \right)d \right]\left[ b'(x)(1-d)+ (N-1)\bar{b}(1-d)+ N\bar{b}\left( 1-c \right)d \right]} \right]$$

Further simplification and using definitions in **Appendix B** results in the expression:

$$\frac{{dw}_{1x}(3)}{d\dot{b}(x)}= d\dot{b}(x)\left( 1-d \right)\left[ \frac{\left[ (1-p\left( x \right)+(N-1)\bar{p} \right]\left[ \left( N-1 \right)\bar{F}\left( 1-d \right) \right]}{\left[ b\left( x \right)\left( 1-d \right)+(N-1)\bar{b}(1-d)+ N\bar{b}\left( 1-c \right)d \right]\left[ b'(x)(1-d)+ (N-1)\bar{b}(1-d)+ N\bar{b}\left( 1-c \right)d \right]} \right]$$

which can finally be simplified to:

$$\frac{{dw}_{1x}(3)}{d\dot{b}(x)}=- d\dot{b}(x)\left[ \left( 1-d \right)g(x)\dot{k}(x) \right]$$

**[D6]**

These offspring are related to the focal individual by $\hat{r}(x)$. Equations **[D2]**, **[D4]**, and **[D6]** can then be summed to give **[B10]** in **Appendix B**.

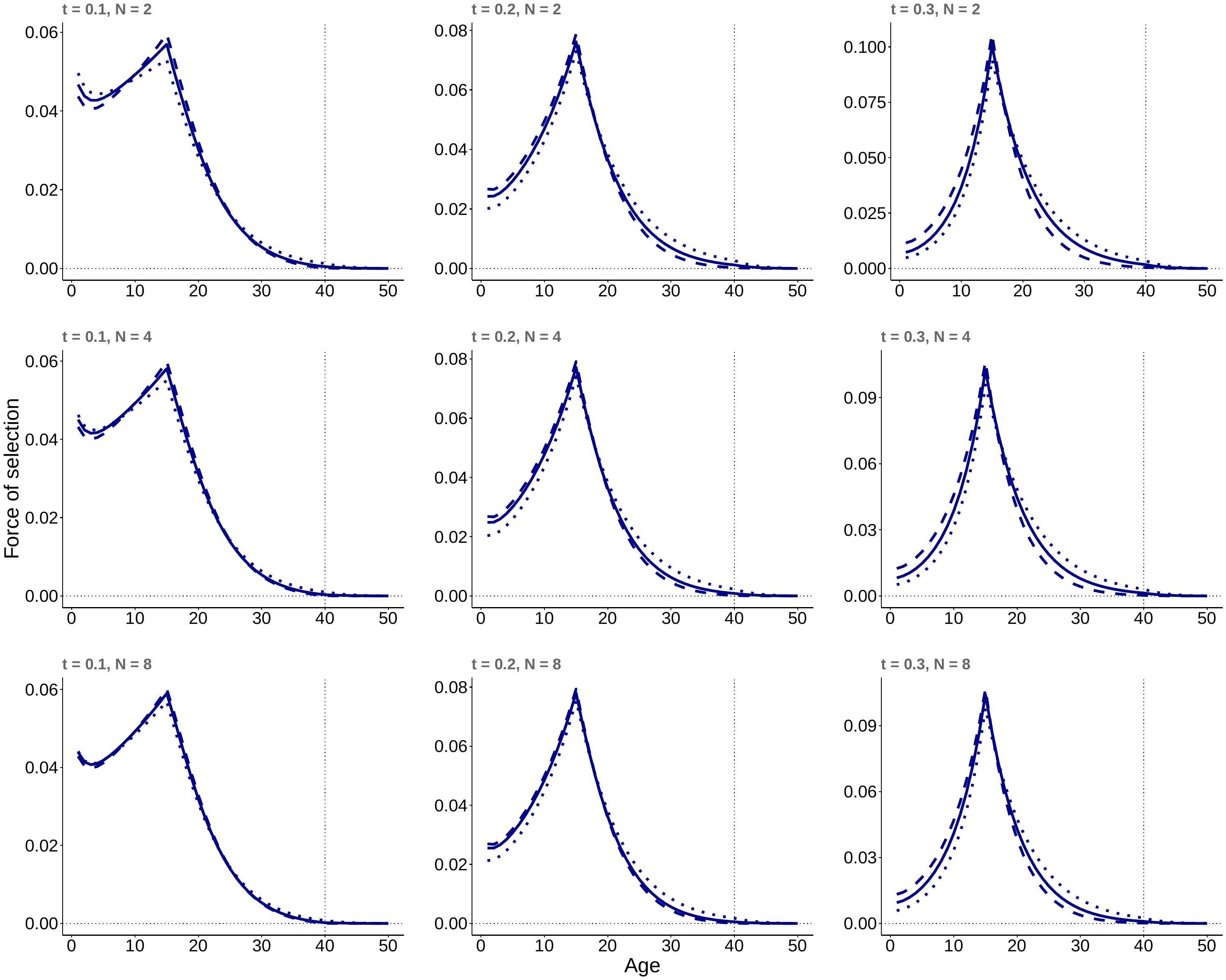

**Figure S1.** The force of selection acting on survival rate at age $x$ in a population with post-reproductive survival ($\mathbf{see Fig.1}$) as a function of the *(i)* the magnitude of the social contribution from post-reproductive individuals to the survival of juveniles, where t = the fraction of juvenile survival that is due to post-reproductive individuals, *(ii)* the number of individuals on the patch (N) and the juvenile dispersal rate in the population: 0.2 (dotted line), 0.5 (continuous line), and 0.8 (broken line). The vertical dotted line at age 40 represents the age at which reproduction ceases.

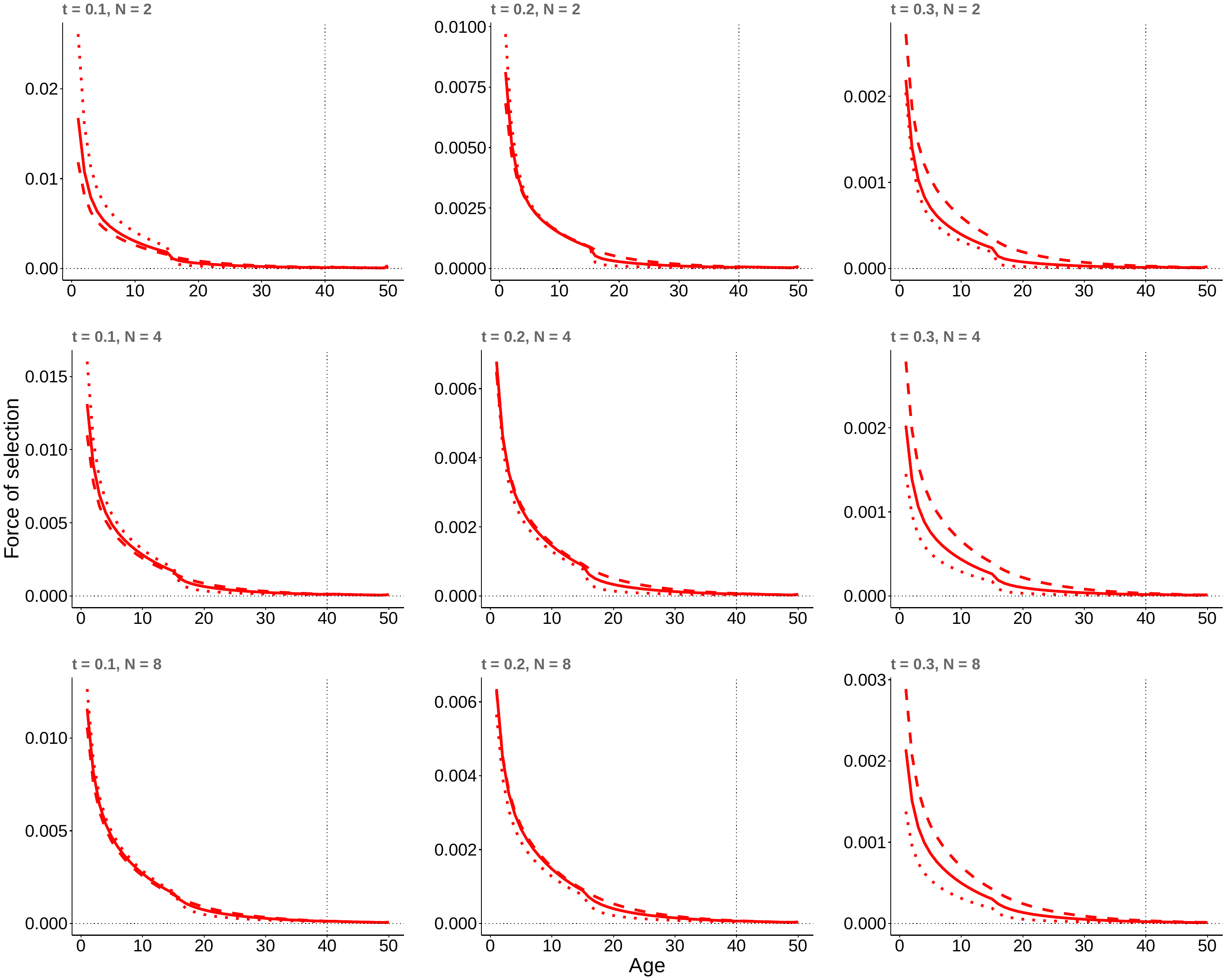

**Figure S2.** The force of selection acting on reproduction at age $x$ in a population with post-reproductive survival ($\mathbf{see Fig.1}$) as a function of the *(i)* the magnitude of the social contribution from post-reproductive individuals to the survival of juveniles, where t = the fraction of juvenile survival that is due to post-reproductive individuals, *(ii)* the number of individuals on the patch (N) and the juvenile dispersal rate in the population: 0.2 (dotted line), 0.5 (continuous line), and 0.8 (broken line). The vertical dotted line at age 40 represents the age at which reproduction ceases.

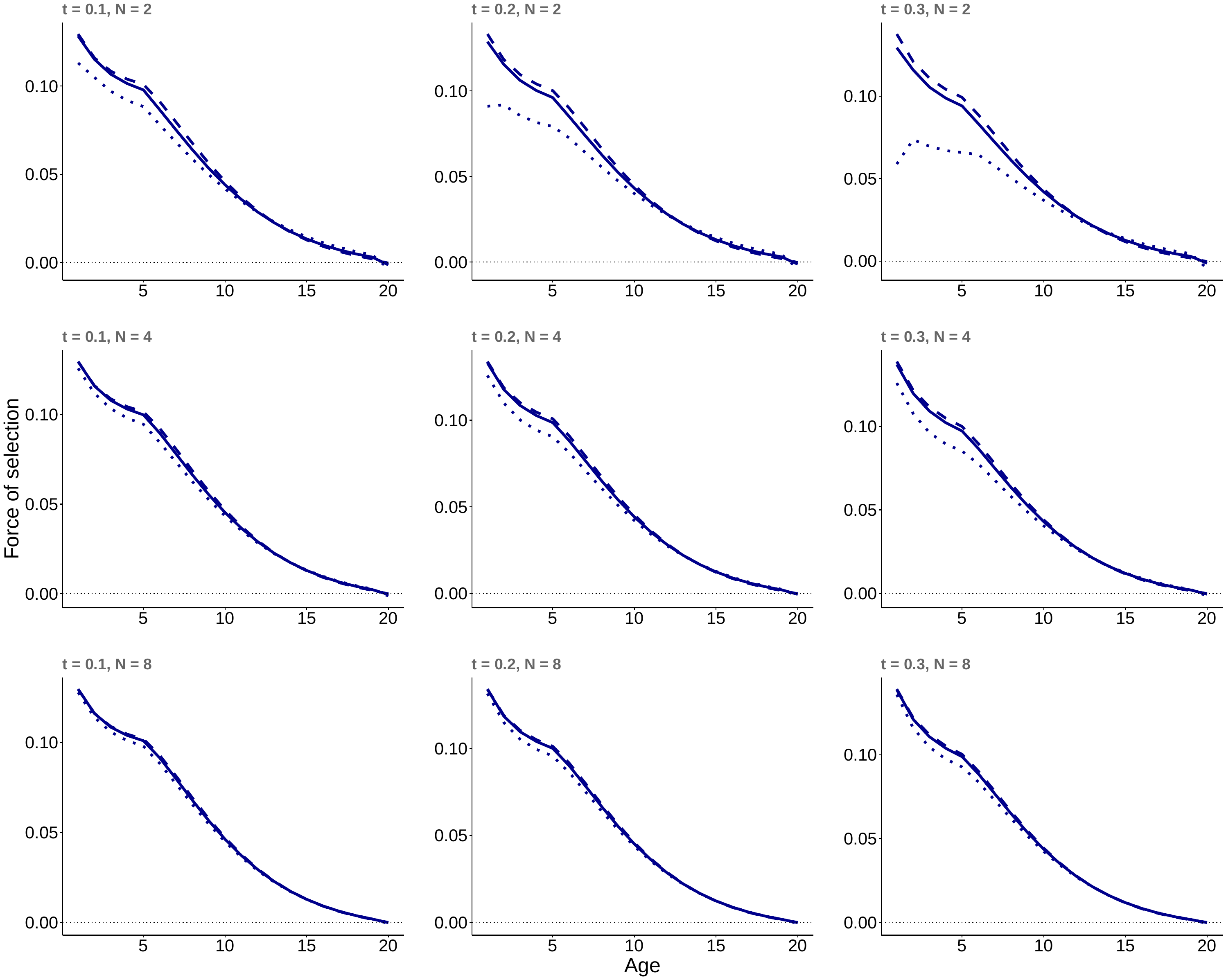

**Figure S3.** The force of selection acting on survival at age $x$ in a population with pre-reproductive helpers $(\mathbf{see Fig.2})$as a function of the *(i)* the magnitude of the social contribution from pre-reproductive individuals to the reproduction of reproductive-aged adults, where t = the fraction of reproduction that is due to pre-reproductive individuals, *(ii)* the number of individuals on the patch (N) and the juvenile dispersal rate in the population: 0.2 (dotted line), 0.5 (continuous line), and 0.8 (broken line).

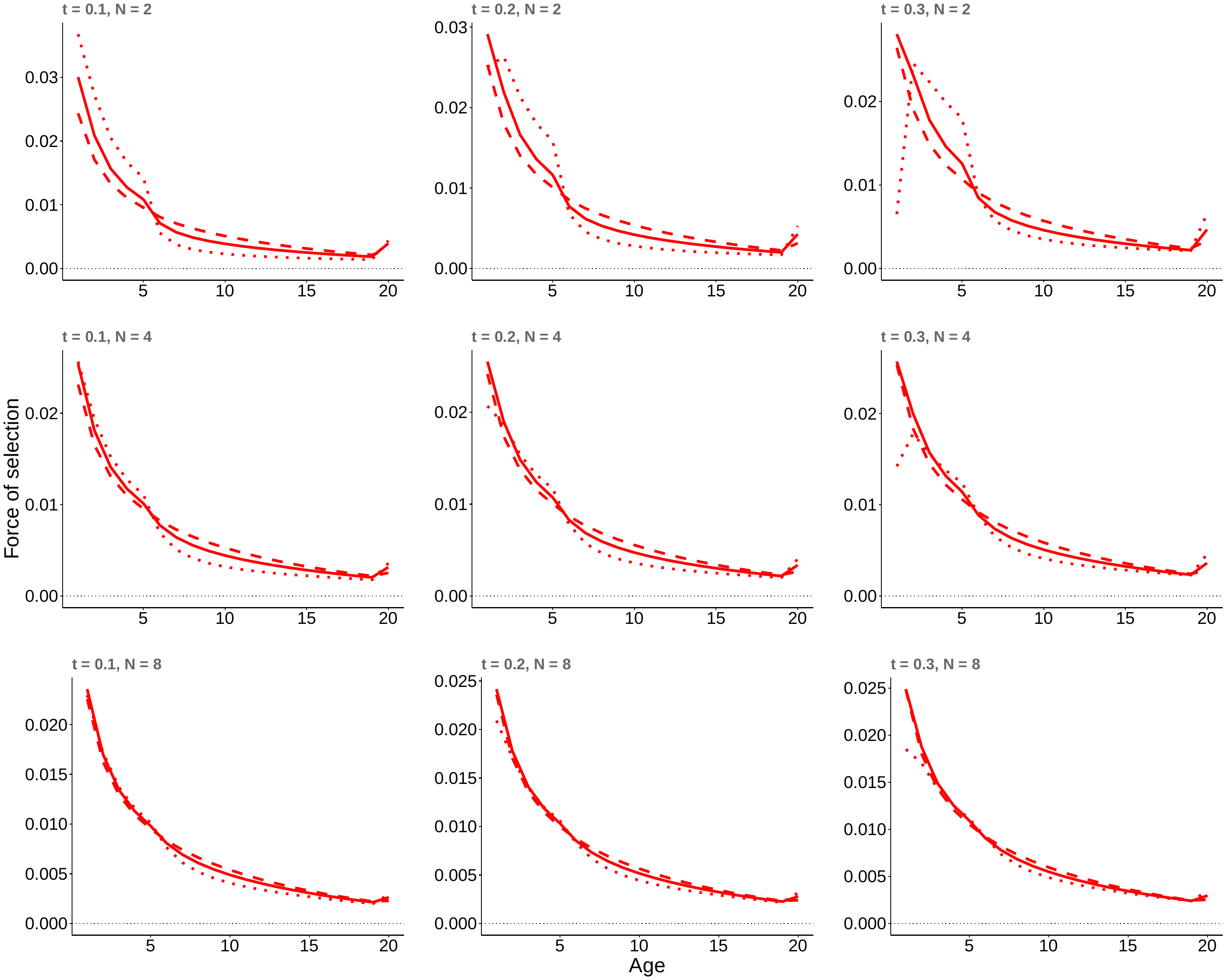

**Figure S4.** The force of selection acting on reproduction at age $x$ in a population with pre-reproductive helpers $(\mathbf{see Fig.2})$as a function of the *(i)* the magnitude of the social contributions from pre-reproductive individuals to the reproduction of reproductive-aged adults, where t = the fraction of reproduction that is due to pre-reproductive individuals, *(ii)* the number of individuals on the patch (N) and the juvenile dispersal rate in the population: 0.2 (dotted line), 0.5 (continuous line), and 0.8 (broken line).
